# Variable phylosymbiosis and cophylogeny patterns in wild fish gut microbiota of a large subtropical river

**DOI:** 10.1101/2024.11.21.624731

**Authors:** Yaqiu Liu, Xinhui Liu, Konstantinos Ar. Kormas, Yuefei Li, Huifeng Li, Jie Li

**Author notes:** Corresponding author: Jie Li, Pearl River Fisheries Research Institute, Chinese Academy of Fishery Sciences, Guangzhou 510380, China.

## Abstract

The persistence and specificity of fish host-microbial interaction during evolution is an important part of exploring the host-microbial symbiosis mechanism. However, it remains unclear how the environmental and host factors shape fish host-microbe symbiotic relationship in subtropical rivers with complex natural environments. Freshwater fish are important consumers in rivers and lakes and considered keystone species in maintaining the stability of food webs there. In this study, patterns and mechanisms shaping gut microbiota community in 42 fish species from the Pearl River, in the subtropical zone of China were investigated. The results showed that fish host specificity is key driver of gut microbiota evolution and diversification. Different taxonomic levels of host showed different degrees of contribution in gut microbiota variation. Geographical location and habitat type were the next most important factors in shaping gut microbiota across the 42 fishes, followed by diet and gut trait. Our results emphasized the contribution of stochastic processes (drift and homogenizing dispersal) in the gut microbial community assembly of freshwater fishes in the middle and lower reach of the Pearl River. Phylosymbiosis is evident at both global and local levels, which are jointly shaped by complex factors including ecological or host physiological filtration and evolutionary process. The core microbiota showed co-evolutionary relationships of varying degrees with different taxonomic groups. We speculate that host genetic isolation or habitat variation facilitate the heterogeneous selection (deterministic process), which occurs and results in different host-core bacterium specificity.

## 1 Introduction

The vertebrate gut contains a variety of microorganisms, especially symbiotic bacteria, which together form complex microbial ecosystems (1). The gut microbiota has been recognized as a vital "microbial organ" of animals and is closely related to many feeding characteristics of its host, which further influences population growth, reproduction and trophic niche differentiation (2–5). In recently years, relevant research suggests that the vertebrates’ adaptive capacity does not only depend on their host genome, but also their symbiotic microbiome (6). Fish are the most diverse group of vertebrates on Earth, with more than 34,000 species living in freshwater, saltwater, and the deep ocean, and they are critical to global ecosystems and food supply (3). There are numerous bacterial symbionts in the gut habitat of wild fishes, which contribute to their host growth, development, and health (7, 8). Despite coevolving with microbial symbionts for over 400 million years, wild fish gut microbiota has been relatively remained largely unknown compared to terrestrial taxa, especially in wild freshwater fish.

Freshwater fish are regarded as the dominant consumers in rivers and lakes. Due to their diverse feeding modes, fish significantly enhance the trophic link and nutrient recycling/retention in aquatic habitats. For this, they are often considered keystone species in maintaining the stability of food webs in rivers and lakes. An essential part of fish nutrition is essentially mediated by their gut microbiota which can synthesize host essential amino acids, vitamins and short-chain fatty acids, regulate intestinal epithelial cell permeability, assist hosts in degrading plant polysaccharides and other nutrients, and improve their digestion and absorption efficiency (9, 10). The specialized intestinal mucosal structure and intestinal core microbiota function can enhance fish tolerance to fluctuations in external resources and improve the efficiency of nutrients extracted from various food sources (4, 11, 12). In addition, gut core microorganisms can also provide vital support for hosts to regulate their energy metabolism balance, improve immune function, and resist pathogen invasion (13, 14).

As gut bacterial symbionts have a profound impact on the nutrition and development of their hosts, as well as their overall fitness, it is critical to answer the question of how hosts maintain these benefits by procuring or inheriting these vital symbionts, which is still largely unanswered. In general, during the first developmental stages, most fish hosts are basically sterile, and the gut microorganisms originate from their external environment or prey (15). Hosts are exposed to a common group of prospective microbial invaders, whereas they selectively filter and choose specific microorganisms to establish a symbiotic relationship with them. Microbial community structure in the habitat environment can often be predicted by changes in environmental factors, such as salinity, temperature and other water environmental factors (16). Theoretically, bacteria can be horizontally transmitted between distantly related hosts by sharing habitats and diets, resulting in similar gut microbiota communities among hosts with overlapping diets and habitats (17). However, hosts have autonomously selected their gut microbiota and regulated the diversity and abundance of gut microbiota by the evolution of innate and adaptive immune systems. These characteristics are determined by the host’s own genotype, and the host population will develop a variant immune response due to genetic diversity, which may in turn help microbial colonization and filtration. Nevertheless, in cases of high habitat overlap and finer phylogenetic scales, it remains unclear whether phylosymbiosis (18) and fish host-microbe co-evolution are determined by the relative importance of the microorganisms to the host’s fitness.

Some studies proposed that the host’s selection of microorganisms is believed to be the major factor responsible for shaping the gut microbial symbiosis (19). At present, it is generally accepted that the gut microbiota assembly is predominantly shaped and regulated by deterministic and stochastic processes. According to the niche theory, the persistence and distribution of species can be significantly influenced by deterministic processes, including biological factors (e.g., biological interactions) and abiotic factors (e.g., environmental filtration) (20). Conversely, neutral theory presupposes that all individuals are ecologically equivalent and that species dynamics and patterns are primarily influenced by stochastic processes, such as migration, speciation/extinction, and random birth/death (21). Deciphering the major elements of microbial community assembly, has crucial theoretical implications for future research into particular host-microbial symbiosis relationship.

The Pearl River is the second largest river in China, located in tropical and subtropical areas, holding over 600 fish species described to date. Abundant and various fish species is hypothesized to be crucial links to its diverse climatic conditions and complex landform structure. Simultaneously, the middle and lower reaches of the Pearl River are the most densely populated and economically active regions. These regions are significantly impacted by human activities, especially the construction of water conservation projects. Such projects can change the environmental factors of river ecosystem like water temperature, eutrophication degree, aquatic microorganisms, dissolved oxygen, stratification etc. and, thus, significantly alter the fishes’ habitat. However, it remains unclear how the environmental and host factors shape fish host-microbe symbiotic relationship in this large and complex subtropical river.

In the current study, we identified gut microbiota of 42 wild fish species belonging to five taxonomic orders (Cypriniformes, Siluriformes, Perciformes, Synbranchiformes, Clupeiformes) in the middle and low reach of the Pearl River by bacterial 16S rRNA metabarcoding. For half of these species, no gut microbiota exist to date. We aimed to (i) uncover the impact of phylogenetic, environmental, and biological factors in shaping the fish gut microbiota; (ii) delineate information on the ecological processes assembling fish gut microbiota; (iii) explore patterns of phylosymbiosis and co-phylogeny for the core gut microbiota of the 42 investigates fishes.

## 2 Material and Methods

### 2.1 Sampling collection

During July and September 2023, 199 wild-caught fish belonging to 42 species in 14 taxonomic families were sampled at 11 locations throughout the Pearl River’s middle and lower reaches (Figure 1A; Table S1). With the help of local fisherman, intact fish using gillnets, cages, or longlines were captured. To determine the influence of diverse phylogenetic and environmental variables on gut microbiota composition, fish species were collected strategically with varying amounts of habitat utilization, gut type, and nutritional preferences. MS 222 (3-aminobenzoic acid ethyl ester methane sulfonate, Sigma, Germany) was used to anesthetize all specimens, after which they were swiftly beheaded. To avoid contamination from the skin surface, the fish body and instruments were pre-sterilized. To limit changes in gut microbiota composition, the average period from extraction to storage was <10 minutes per fish. Each sample, consisted of the gut tissue with its contents, was immediately stored in liquid nitrogen. Upon return to the laboratory, all samples were transferred swiftly to a -80℃ ultra-low refrigerator until further experiments.

**Figure 1.**
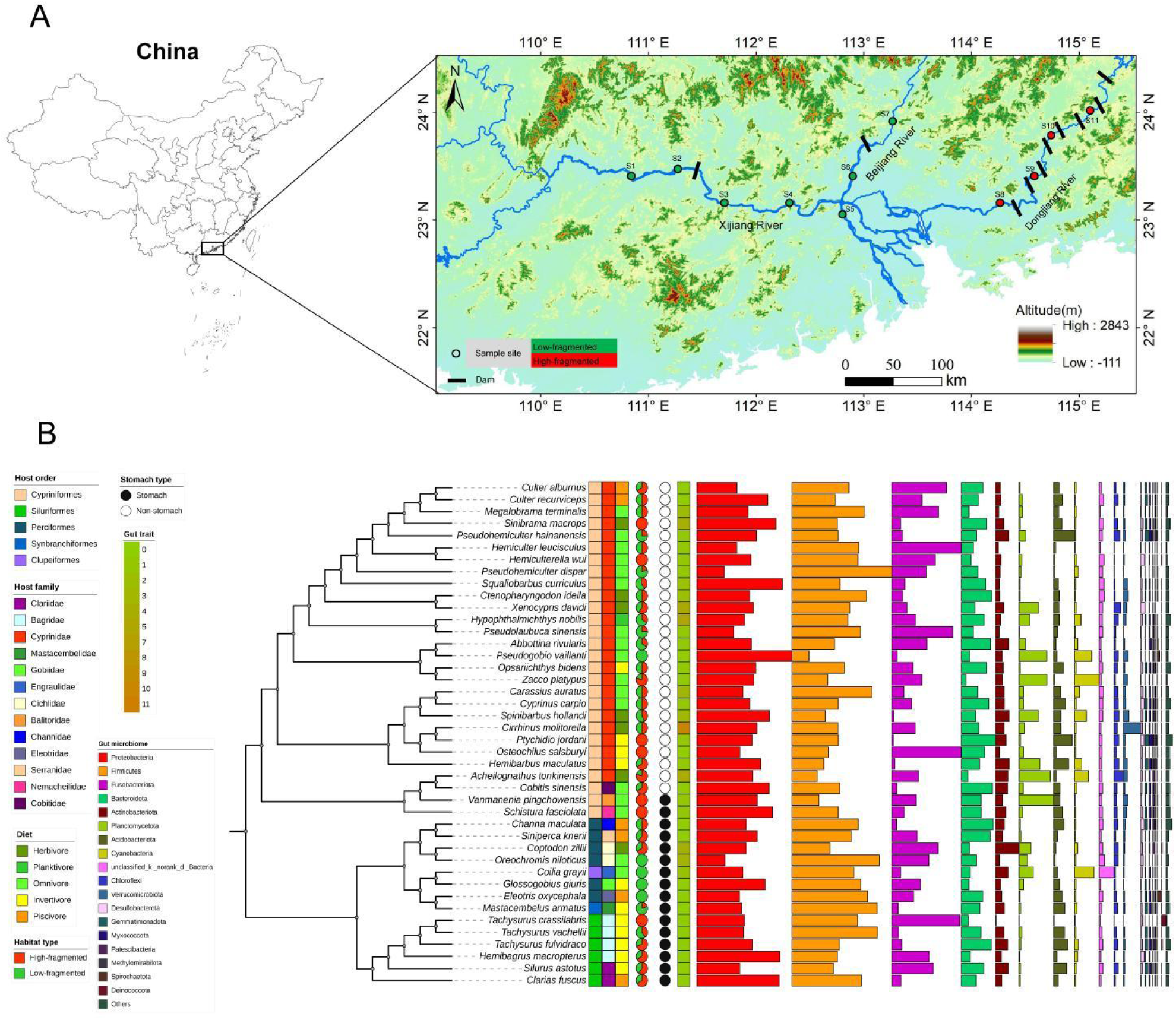
Broad microbiome in the fish gut in the middle and lower research of the Pearl River. (A) Map showing the geographical locations of fish samples analyzed in this study. (B) Relative abundance of bacterial taxa in the gut microbiome in fish host species. Microbial community compositions are displayed based on the phylum level. Host phylogeny was inferred from COI gene evolution history using iq-tree. For convenience, the lengths of branches do not represent evolutionary distance. Host metadata are labelled using different colours and shapes.

The taxonomic information (family, genus, and species) of all the samples was determined by examining their morphological features as described in the literature (22–24). For each specimen, body size (length and weight) and sex were recorded. Gut trait was characterized using relative gut length (gut length/body length). Relative gut length was used to reduce the effect of individual differences on gut length. Dietary habits exhibited significant variation as reported in earlier research (25) and was determined according to FishBase (https://fishbase.net.br/search.php). These feeding habits included piscivores, herbivores, invertivores, omnivores, and planktivores (Table S1). In each sampling site, water temperature, dissolved oxygen (DO), total dissolvable solid (TDS) conductivity and pH were assessed using an HQ30 equipment (Hach Company in Loveland, CO, USA). The transparency (secchi depth, SD) of water was measured on site with Sachter’s plate. Water sample was filtered by WHATMAN GF/C filter membrane, the content of chlorophyllin a (Chla) in water body was determined by spectrophotometry. Water samples were obtained at each sampling station, namely 0.5 meters below the water surface, in order to analyze the levels of nitrogen (TN) and phosphorus (TP). The levels of turbidity, total nitrogen (TN), and total phosphorus (TP) were measured by analyzing three identical samples taken from each water sampling location (26). All the environment parameters are presented in Table S1.

### 2.2 DNA extraction, amplification, and high-throughput sequencing

Total genomic DNA was isolated from 2 grams of intestinal contents using the QIAamp DNA Stool Mini kit (Qiagen, Valencia, CA), following the manufacturer’s stated methodology. The hypervariable V4 region (515F–806R) was targeted for amplification of the 16S rRNA genes from different locations, as described by Caporaso et al. (27). A 1% agarose gel extract was used to purify the polymerase chain reaction (PCR) product and quantify it using the Qubit 4.0 (Thermo Fisher Scientific, Waltham, CA) according to the manufacturer’s instructions. The amplicons were combined in equal amounts and subjected to sequencing using the Illumina PE250 platform (Illumina, San Diego, CA, USA) following standard procedures provided by Majorbio Bio-Pharm Technology Co., Ltd. Shanghai, China.

### 2.3 Sequence data processing

The 16S rRNA gene sequences were analyzed using the QIIME2 environment (28). At first, the raw data were grouped according to their barcode sequences. Subsequently, the primers and chimeric sequences were removed using Cutadapt and DADA2, respectively (29, 30). A Naïve Bayes classifier was trained using the SILVA (version 138) 16S reference sequences for the purpose of assigning sequences into taxonomic groups (31). Sequences that were not classified, together with those labeled as ’mitochondria’, ’chloroplast’, ’archaea’, and sequences with taxonomic precision restricted to the domain level, were excluded from further investigations. Sequences occurring as singletons and doubletons in the whole dataset were also excluded, as they are likely errors of the sequencing process.

### 2.4 Host phylogenetic tree reconstruction

The cytochrome c oxidase subunit I (COI) genes linked with the present fish samples were retrieved from the NCBI database (https://www.ncbi.nlm.nih.gov/) based on their species information. The COI sequences were first aligned using mafft version 7.310, as described by Katoh and Standley in 2013. Subsequently, the alignment was fine-tuned using trimal version 1.4.rev15 and configured with the -fasta option (32). Model finder was used to discover the best acceptable model (33). The building of maximum-likelihood trees was conducted using iq-tree version 1.6.11 (34) with 1000 bootstrap replications carried out using ufboot2 (35). The trees were shown in ITOL version 6.

### 2.5 Bioinformatics analysis

After elimination of the undesirable sequences, alpha-diversity metrics (Faith’s PD index) were computed in QIIME2. Rarefaction plots were produced to examine if the sequencing efforts were adequate to encompass the variety of the bacterial community. The Kruskal-Wallis (KW) test was used to analyze the variations between groups, and the statistical significance was assessed by adjusting the p-value using the Bonferroni correction. The vegan R package was used to generate Principal Coordinates Analysis (PCoA) plots, which visually represent the similarities in microbiota across samples based on the Bray-Curtis distance matrix. The Binary-Jaccard and Bray-Curtis dissimilarity distance matrices were used to determine beta-diversity (pairwise distances) across groups. This calculation was performed using the ordinate function in the vegan R package (36). Permutational multivariate analysis of variance (PERMANOVA) tests were conducted to examine the impact of environmental and host-related variables on the beta-diversity distances of bacterial community composition. The impacts of these factors were assessed using adonis (PERMANOVA) analysis. The statistical significance scores for both tests were computed using 999 permutations. β-dispersion was assessed using the betadisper function in the vegan R package. This was done by calculating the non-euclidean distance between each sample and the group centroid at various host taxonomic levels. A higher β-dispersion value indicates a greater difference in the composition of the gut microbial community within the group. We defined the core bacterial taxa of fish as those with presence in ≥ 90% of all samples and ≥ 95% occurrence in each fish orders. ParaFit and PACo (Procrustean Approach to Cophylogeny), was used to assess the co-evolutionary relationship between the host and their symbionts (core genus level) by testing their signal of codiversification (37, 38). The signal of cophylogeny/codiversification was independently verified via the implementation of three distinct approaches. The statistical significance values for all three tests were computed using 999 permutations. Subsequently, the phylogenetic signals of ASVs and the local indicator of phylogenetic association (LIPA, local Moran’s I) were computed using the phylosignal R package with 9999 permutations (39) .

### 2.6 Microbial community assembly and stochasticity

The relative value of community assembly methods was examined according to the technique of Stegen et al (40). The null model, consisting of 1000 randomizations, was used to compute the β-nearest taxon index (βNTI) and the Raup-Crick index based on Bray-Curtis dissimilarity (RCbray). βNTI values less than -2 and βNTI values greater than 2 were viewed as indicating homogenous selection and variable selection, respectively. Both of these processes are deterministic.

|βNTI| < 2 and RCbray< −0.95 indicate homogeneous dispersal, |βNTI| < 2 and RCbray>0.95 represent dispersal limitation, and |βNTI| < 2 and |RCbray| < 0.95 suggest an undominated process (also referred to as drift), all of which are stochastic processes (21).

## 3 Results

### 3.1 Gut bacterial composition

A total of 15.54 million Illumina sequences from the 16S rRNA gene’s hypervariable V4 region were retrieved. The rarefaction curves reached an asymptote for the majority of the samples, indicating that the sequencing effort was adequate in capturing the microbial diversity of the examined fish (Figure S1A). Good’s coverage of the samples was 0.991 ± 0.003, further supporting this conclusion (Figure S1B). The taxonomic categorization of bacterial sequences identified 548 and 1022 bacterial species at the order and family levels, respectively. Proteobacteria were the top-ranked group, accounting for a relative abundance ranging from 13.8% to 47.5%, followed by the that the Firmicutes (8.4%-49.9%), Fusobacteria (0.83-34.8%) and Bacteroidota (2.3-17.1%) (Figure 1B). Collectively, these four phyla represented over 50% of the overall sequences found in all fish species. *Cetobacterium* was the most abundant bacterial genus, comprising 11.75% of the total abundance. Additional abundant bacterial genus were *Clostridium_sensu_stricto_1*, *Romboutsia*, *Aeromonas*, *Lactobacillus*, *Bacteroides* (Figure S2). There was a positive correlation between relative abundance and prevalence. A total of eight core bacterial genera were observed in the studied species (Figure 2) in the following decreasing order of relative abundance: *Cetobacterium, Clostridium*_sensu_stricto_1, *Aeromonas*, *Romboutsia*, *Bacteroides*, *Lactobacillus*, *Achromobacter*, and *Bacillus* (Figure 2). The predominance of these genera was observed across the marsupial orders Cypriniformes (≥90.23%), Perciformes (≥96.15%), and Siluriformes (≥86.67%). In addition to their high prevalence, these eight genera were also abundant across the three orders (Figure S3).

**Figure 2.**
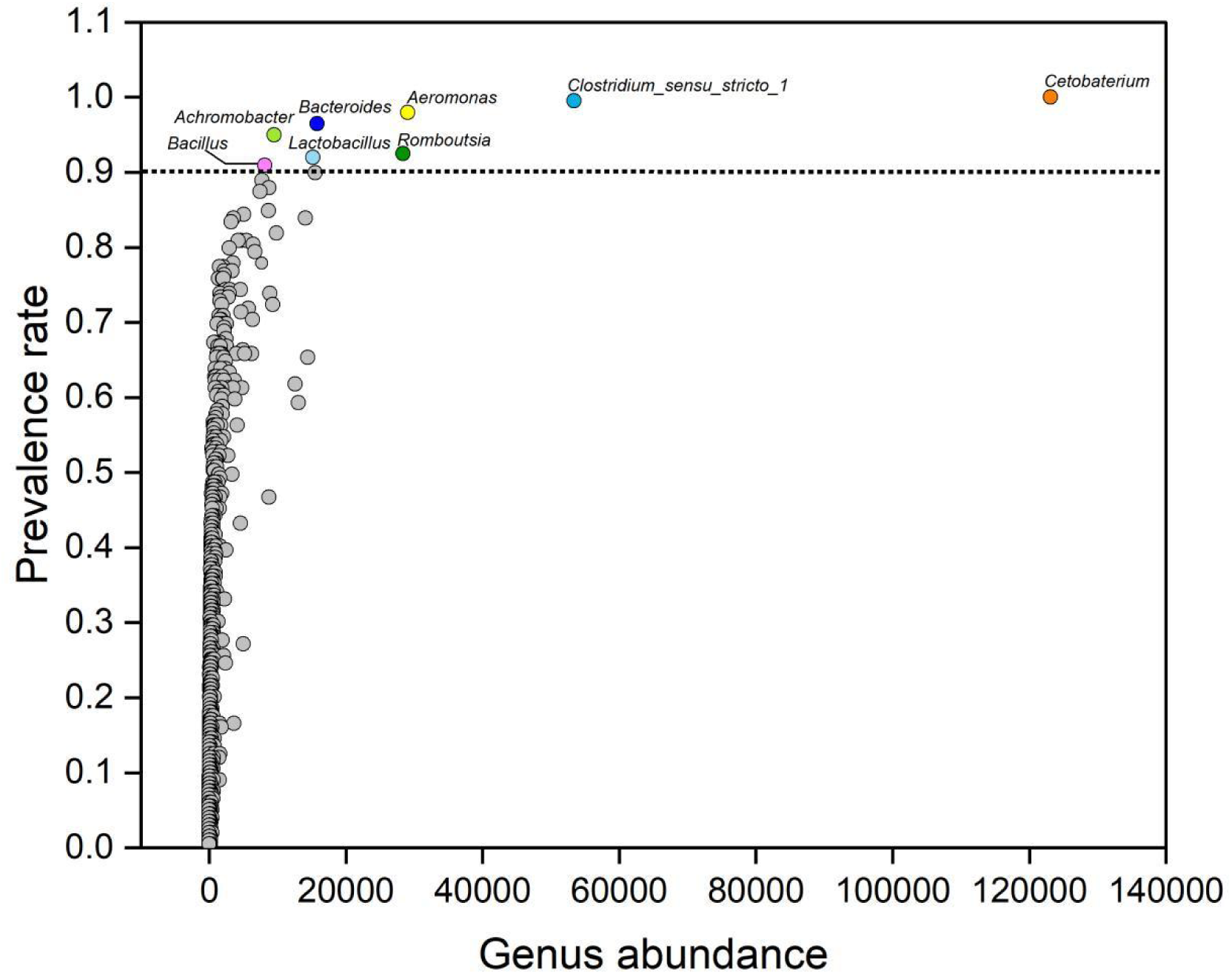
Abundance and prevalence of bacterial genera in all fish host. Prevalent rate of genus of all samples≥ 90% was highlighted in the different colors, and other genus ≤ 90% was shown in gray colour.

### 3.2 Host factors shape gut microbiota diversity and community structure

The study assessed the impact of several factors on the alpha diversity (i.e., the Faith’s PD) of bacterial communities in fish gut (Figure S4). These factors included host species, geographical location, diet, stomach type, habitat type, and sex. The Faith’s PD values of exhibited substantial variation between gut bacterial communities from different species (Kruskal-Wallis H test *P*< 0.05), geographical location (Kruskal-Wallis H test *P*< 0.01), and habitat types (Wilcoxon rank-sum *P*< 0.001). The result of PERMANOVA analysis showed that the significant differences between samples were influenced by the assessed phylogenetic, biological, and environmental variables to different extents (Table 1). The host species accounted for the highest proportion of overall variation in Bray-Curtis (*R*^2^= 33.6%), and Binary-Jaccard (*R*^2^= 25.9%). We also found that the genus (*R*^2^= 22.7%– 29.6%), family (*R*^2^= 6.9%– 9.3%), order (*R*^2^= 2.6%– 4.3%) identity of the samples explained gradually declining proportion of the beta distance variations, suggesting that the bacterial assemblage was gradually less predictable as phylogenetic scope widened (Table 1). Similarly, PCoA plots revealed that samples from the same genus and species tended to cluster together (Figure S5).

**Table 1.** The results of PERMANOVA (Adions) showing the contribution of different phylogenetic, environmental, and biological factors to the between sample variability of the fish gut microbiome in the middle and lower research of the Pearl River.

| Main factors | Bray-Curtis |  |  | Binary-Jaccard |  |  |
| --- | --- | --- | --- | --- | --- | --- |
| | F.Models | $R^2$ | $P$-Value | F.Models | $R^2$ | $P$-Value |
| Host order | 2.155 | 0.043 | 0.001 | 1.283 | 0.026 | 0.004 |
| Host family | 1.451 | 0.093 | 0.005 | 1.053 | 0.069 | 0.250 |
| Host genus | 1.828 | 0.296 | 0.001 | 1.277 | 0.227 | 0.001 |
| Host species | 1.938 | 0.336 | 0.001 | 1.339 | 0.259 | 0.001 |
| Geographical location | 2.505 | 0.118 | 0.001 | 1.640 | 0.080 | 0.001 |
| Habitat type | 12.975 | 0.062 | 0.001 | 4.971 | 0.035 | 0.001 |
| Diet | 1.682 | 0.034 | 0.012 | 1.300 | 0.026 | 0.033 |
| Stomach type | 3.264 | 0.016 | 0.012 | 1.964 | 0.010 | 0.029 |
| Gut trait | 8.631 | 0.081 | 0.001 | 3.246 | 0.032 | 0.001 |
| Sex | 0.650 | 0.003 | 0.750 | 0.853 | 0.004 | 0.636 |
| All factors | 0.496 |  |  | 0.391 |  |  |

Moreover, geographical location and habitat type also play important roles in effecting on the beta-diversity of fish gut microbiota explaining 8.0% – 11.8% and 3.5% – 6.2%, respectively (Table 1). Decomposing all environmental factors and geographical factors by variance partitioning revealed that the variables jointly only explained 1.49% of the variance (Figure S6A). The ecological factors (3.7%) and geographical factors (1.44%) both showed small variance explanatory rate exclusively. We searched for environmental factors that directly influenced fish gut microbiota based on variance partitioning. The environmental factor with the greatest explanatory power was DO, which explained 18.01% of the variance in microbial community diversity (Table S2). To further reconstruct the relationship between environmental parameters and the fish gut bacterial community between high and low fragmented habitat type, we calculated the correlations of Bray-Curtis dissimilarities of community composition and environmental parameters with Euclidian distances using a Mantel test. Overall, DO and TDS were significantly correlated with taxonomic composition in fish gut bacterial communities of low fragmented habitat (Mantel’s *r* = 0.073 and 0.079, *P* < 0.01, Figure S6B). Simultaneously, pH showed strong relationships with taxonomic composition in fish gut bacterial communities of high fragmented habitat (Mantel’s *r* = 0.053, *P*<0.05, Figure S6B). On the other hand, four biological factors including diet (2.6% – 3.4 %), stomach type (1.0% – 1.6%), gut trait (3.2% – 8.1 %), and sex (0.3% – 0.4%) also accounted for some proportion of the bacterial variability (Table 1). Our results showed that host gut trait explained highest proportion of the beta distance variations of different samples. In addition, our results indicated that there was a positive linear relationship between the relative abundance of Planctomycetota, Chloroflexi, and Verrucomicrobiota and host relative gut length, whereas a negative correlation between the relative abundance of Bacteroidota and the host relative gut length was observed (Figure S7).

### 3.3 Ecological processes affecting bacterial community assembly

We quantified the impact of specific deterministic (homogenous and heterogenous selection) and stochastic (homogenizing dispersal, dispersal limitation, and drift) processes on the bacterial community assembly (Figure 3 and S8). The results showed that the contribution of stochastic (dispersal limitation and drift) processes in shaping bacterial community in all host orders, except the Clupeiformes(Figure 3A). Deterministic (homogenous and heterogenous selection) processes seem to drive gut microbiota assembly of the Clupeiformes (Figure 3A). Furthermore, our results showed that homogenous selection is the sole deterministic process that shapes community assembly across mostly families, whereas another deterministic (heterogenous selection) process contributed in assembling microbial community in samples of the Engraulidae (24.5%), Cyprinidae (3.0%), and Bagrideae (1.4%) (Figure 3B). Drift (20%–77.7%) was the most significant factor in the formation of community assembly within stochastic processes, accounting for all families. Homogenizing dispersal had a negligible impact. In addition, the relative contribution of ecological processes varied across datasets associated with various diet, stomach types, and habitat (Figure 3C, D, E), while relative contribution of ecological processes was little discrepancy in different gender groups (Figure 3F). The relative importance of dispersal limitation in shaping microbial community of fish host in the low fragmented habitat was much lower than that in the high fragmented habitat. We further explored the relationships between the βNTI and microbial Bray–Curtis similarity were used to infer the impact of deterministic/stochastic assembly processes on the fish gut bacterial community. The result indicated that gut microbiota similarity had a negative correlation with βNTI (Figure 3F).

**Figure 3.**
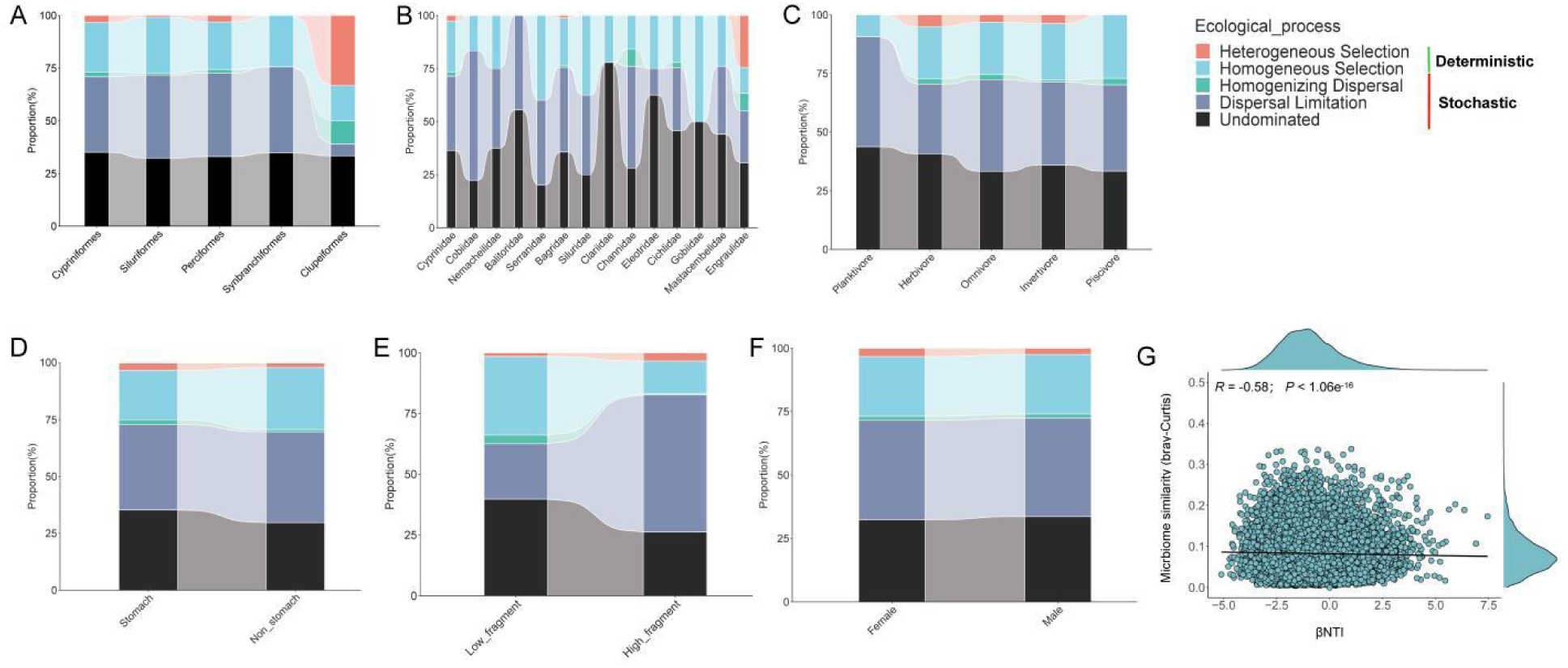
Ecological processes about the gut microbial community assembly. Quantification of deterministic and stochastic processes governing the microbial community assembly and the percentages are relative contributions of each process turnover for different orders (A), families (B), diet (C), stomach types (D), habitat (E), and sex (F). (G) the relationship between βNTI and the microbiome similarity (Bray-Curtis distance). Linear regression models (shown as red lines) and associated correlation coefficients are provided on each panel.

### 3.4 Evidence of phylosymbiosis in fish gut microbiota

In order to investigate any phylosymbiosis patterns in the investigated fish holobionts, we firstly calculated host genetic similarity and microbial dissimilarity (ASV level) by using the COI gene sequence and Bray-Curtis distance, respectively. It was shown that host COI gene similarity and gut microbial dissimilarity had a significant positive correlation (Figure 4A; *R* = 0.21, *P* < 2.2e^-16^). The UPGMA clustering tree indicated the high bacterial community (genus level) similarity between different host species based on Bray-Curtis distance matrix using a hierarchical clustering method (Figure S2). The result confirmed that the similarity of gut microbial communities of closely related species in Cypriniformes. Subsequently, we quantified the microbial community composition heterogeneity at various host taxonomic levels (order down to species) using β-dispersion analysis (Figure 4B). The result showed that host samples classified at the species level were the lowest among the four distinct distance matrices, suggesting that the composition of the gut microbial community was closely correlated with the host species level (Figure 4B). A Venn diagram was employed to construct and visualize the shared and unique ASV in gut samples from various orders. We observed 18 prevalent ASVs occuring in all orders, belonging to the *Cetobacterium*, *Aeromonas*, *Clostridium*_sensu_stricto_1, *Romboutsia*, *Achromopacter*, *Pseudomonas*, *Lactobacillus* (Figure 4C).

**Figure 4.**
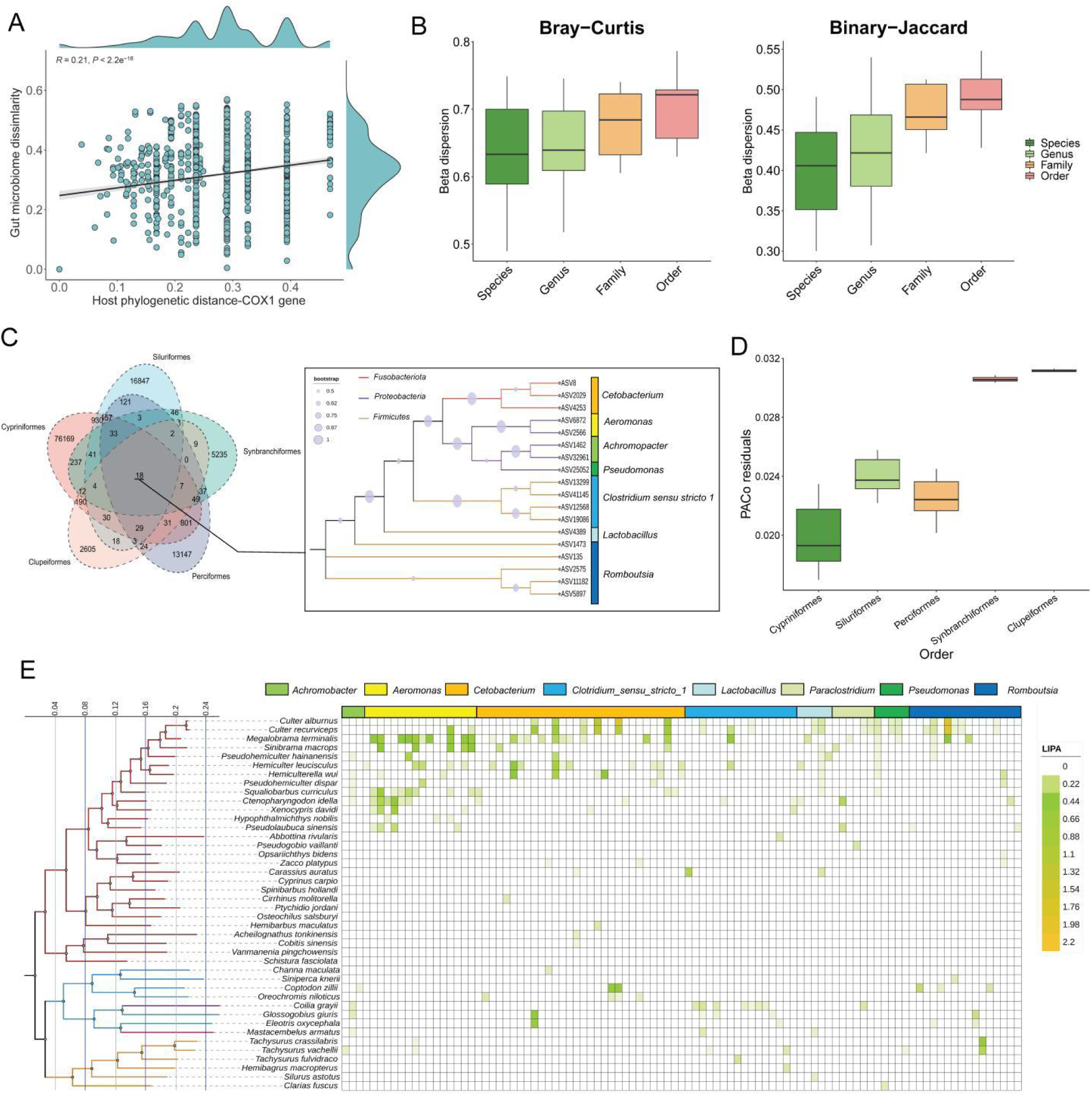
Phylosymbiosis patterns of fish gut microbiome in the middle and lower research of the Pearl River. (A) Linear regression analysis with the slope of the regression line for host COI gene similarity versus microbial dissimilarity. (B) β-dispersion analysis indicating microbial community composition heterogeneity at different host taxonomic levels. (C) Sharing and specificity ASVs of fish gut microbiota from different orders. Neighbor-joining phylogenetic tree of common ASVs. (D) Procrustean approach to co-phylogeny (PACo) analysis between fish hosts and microbiome. (E) Local indicator of phylogenetic association (LIPA) values for correlations between abundances of ASVs (different genera) and specific hosts using heatmap. The host species tree on the left were found to have significant associations with at least one ASV of microbiome (*P*-value < 0.05). Branch colours are determined by host order in the phylogenetic tree.

The PACo results showed host-microbiota residuals of Cypriniformes was lowest, followed by Perciformes and Siluriformes, and host-microbiota residuals of Clupeiformes was highest (Figure 4D). Four of the eight core bacterial genera, including *Cetobacterium*, *Clostridium*_sensu_stricto_1, *Romboutsia,* and *Bacteroides* showed statistically significant cophylogeny signals in two cophylogeny tests conducted when all fish species were analysed together (Table 2). Subsequently, we choose three host orders, including Cypriniformes, Perciformes, and Siluriformes (number of species over three), for cophylogeny testing with these four core genera (*Cetobacterium*, *Clostridium*_sensu_stricto_1, *Romboutsia,* and *Bacteroides*) separately. Our results showed that Cypriniformes and *Cetobacterium, Clostridium*_sensu_stricto_1, and *Romboutsia* showed cophylogeny (Table 2). However, no cophylogenetic signals were observed between Siluriformes and four core genera. Notably, *Cetobacterium* showed strong co-phylogenetic signals to Perciformes, implying that it may play an important role in Perciformes ecology and evolution. Additionally, we calculated local indicator of phylogenetic association (LIPA) values for investigating the potential phylogenetic signals between host phylogenetic tree and gut microbiota (Figure 4E). In the host tree, 97 ASVs were identified with substantial local phylogenetic signals *Cetobacterium* (33 ASVs), *Clostridium*_sensu_stricto_1 (16ASVs) were the most prevalent among the eight genera into which the LIPA-ASVs were classified. *Romboutsia* (14 ASVs), *Aeromonas* (12 ASVs), *Paraclostridium* (six ASVs), *Lactobacillus* (five ASVs), *Pseudomonas* (five ASVs), and *Achromobacter* (three ASVs) comprised the remaining six genera. These eight genera belonged to Fusobacteriota, Protebacteria, and Firmicutes. This illustrates the significant variation in the distribution of LIPA-ASVs among their respective host lineages (Figure 4E). The average number of LIPA-ASVs in each Cypriniformes host species was higher than that of species from other fish orders. Distribution of LIPA-ASVswas concentrated in Cultrinae (Cyprinidae; Cypriniformes). Further, LIPA-ASVs (*Cetobacterium*and *Romboutsia*) were widely distributed in *Culter alburnus*, *Culter recurviceps*, and *Megalobrama terminalis* belonging to the Cultrinae. Notably, we found that abundant distribution of LIPA-ASVs (*Aeromonas*) in *Cetnopharyngodon idella*, *Squaliobarbus curriculus*, *Sinibrama macrops, M. terminalis and Xenocypris davidi*. In addition, we observed four LIPA-ASVs (*Cetobacterium*) and two LIPA-ASVs (*Romboutsia*) with strong co-phylogenetic signals in the species from Perciformes and Siluriformes separately. Overall, the ASV-specific phylogenetic signal was significant associated with Cyprinidae species, indicating its potential importance in the ecological functions and evolution of Cyprinidae.

**Table 2.**
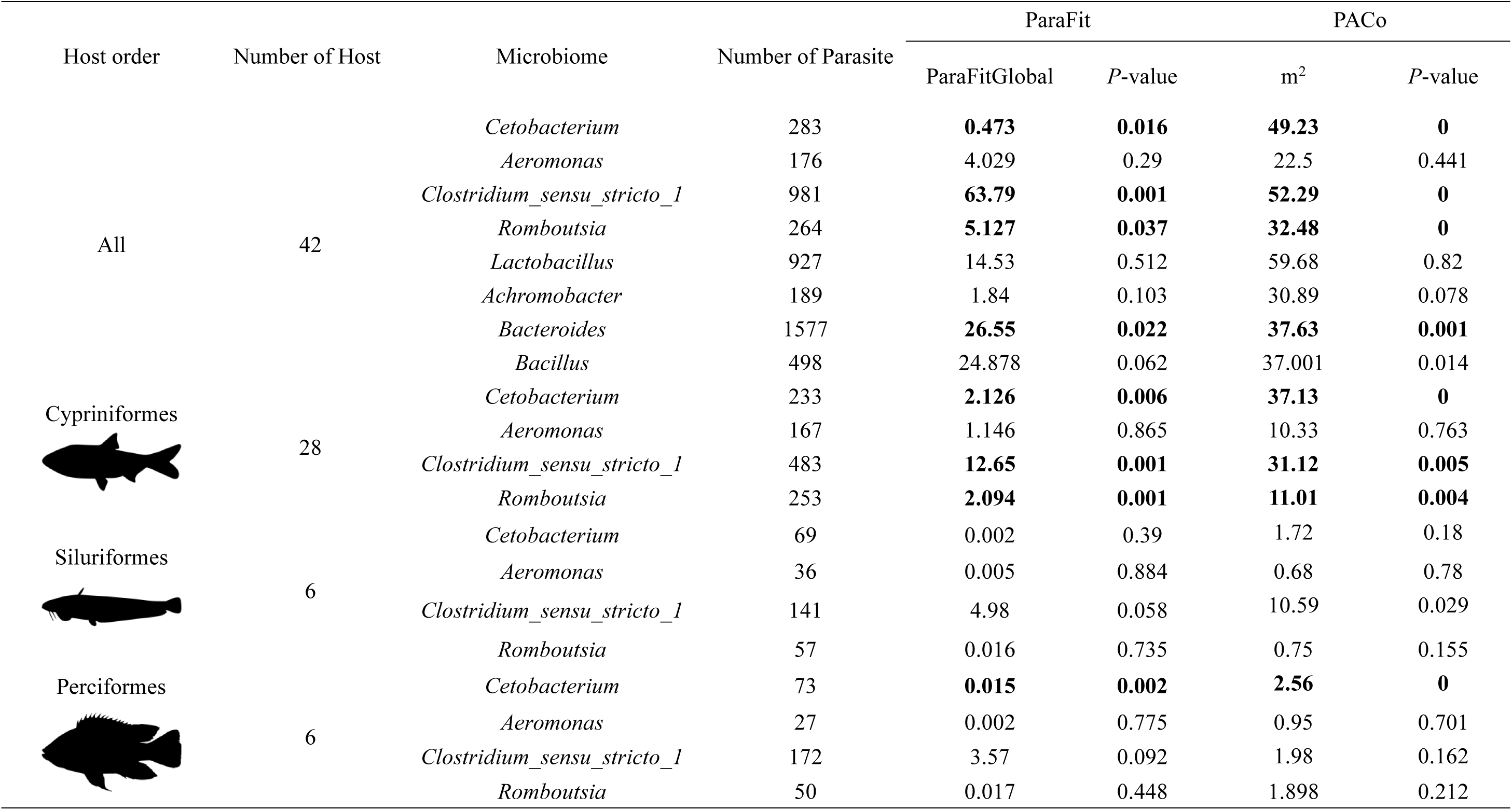
Co-phylogeny associations between fish host and the four core bacterial genera. Test statistics from the tests: ParaFit = ParaFitGlobal; PACo = global goodness-of-fit (m^2^). that showed statistically significant signals of codiversification (P < 0.05) in at least one out of the two tests.

## 4 Discussion

### 4.1 Factors shaping the gut microbiota of freshwater fish

Uncovering the patterns of gut microbial communities adjusted by different factors is fundamental for improving host physiological performance. Studies have shown that feeding preference, rather than host phylogeny, is considered to be the main driver of gut microbiota diversity in marine fish (9). In addition, habitat environment and geographical distance are also considered to be the main factors affecting the diversity of fish gut microbes (14, 41, 42). However, the observed host-specific patterns can be confounded by sharp differences in the habitat and dietary preferences of the species being studied. Recent research has offered that marine fish host specificity and feeding habits are key drivers of gut microbiota variation (11).

In the present study, we investigated the gut microbial communities of 199 fish individuals from 42 species across 14 families and five orders with diverse dietary and habitat preferences to delineate the effect of phylogenetic, biological and environmental drivers variation of gut microbiota composition. Our findings showed that host taxonomic category (species level) identity is the strongest predictor of the gut microbiota (Table 1). Generally, the higher the classification level, the higher the explanation rate of the composition variability of fish gut microbiota, implying a subset of bacterial lineages in fish species is host specific and determined by genetic factors of the fish host. The gut microbiota of vertebrates is host-specific and arose as a result of co-evolution between hosts and microbes (43). Research of Tsang et al. (17) found similar host-specific impact on the gut microbiota of invertebrates. In addition, the immediate environmental factors of host habitat and host diet also had important impact on the microbial composition. Previous studies showed that the gut microbiota of fish is primarily determined by the fish habitat, rather than by genetic factors, which is not exactly consistent with our results (14). The reason may be the striking difference between marine and freshwater habitats investigated by Kim et al. (14), which can mask phylosymbiosis between the host and the gut microbial system. Notably, we observed that host gut trait (relative gut length) significant influenced its gut microbiota. Likewise, there was also significant relationship between the relative abundance of some bacteria and relative gut length of host (Figure S6). This may be due to the fact that gut traits are affected by fish phylogeny, habitat, and trophic level as well as some symbiotic microorganisms in similar ways (12, 44).

### 4.2 Symbiotic microbial community assemble in gut of freshwater fish

Our results emphasized the contribution of stochastic processes (drift and homogenizing dispersal) to the microbial community assembly of freshwater fish inhabiting the middle and lower reach of the Pearl River (Figure 3). The contribution of these processes was exceeding that of deterministic processes, at least in the fish orders Cypriniformes, Siluriformes, Perciformes, and Synbranchiformes. Nevertheless, the importance of deterministic processes (heterogeneous selection) in Clupeiformes was sharply increased. This phenomenon may be related to the inclusion of only one species (*Coilia grayii*) which was sampled from the order Clupeiformes. More importantly, diversification of migratory activities of *C. grayii* in the Pearl River has been reported, and it has the ability to use both fresh and brackish water in the course of reproductive migration (45). Therefore, its life-history has experienced constant environmental changes in brackish water and freshwater environment, and it has been shown that there is a remarkable difference of fish gut microbiota between freshwater and marine habitats (14). Heterogeneous selection may cause communities to diverge if they undergo exposure to distinct environmental conditions (46). Further findings claimed that the discrepancy in the assembling process of gut microbial community in the different fish families is startlingly apparent (Figure 3B). The cumulative effect of historical randomness factors leads to host genetic differentiation that drives different host-bacterial specificity (43, 46). Moreover, gut microbial assembly in planktivorous species is relatively more subject to dispersal limitation (stochastic process) than fish with other feeding item (Figure 3C). This result suggests that highly specialized diets narrowed the range of food and foraging environments, which leads to dispersal limitation of microbiota (43,47). In addition, the different contribution of ecological process (drift and homogeneous selection) in bacterial community between stomach and non-stomach species (Figure 3D), suggests that host biological factors also had some impact on the gut bacterial communities. Notably, our results showed that the dispersal limitation increased, and homogeneous selection decreased in affecting bacterial community in fish from low to high fragmented habitat (Figure 3E). These findings imply that different environmental selective pressures for fish may have caused significant differences of gut microbial community. Specifically, cascade of water conservation project intensifies river fragmentation and impedes communication between different fish populations, which further limit the dispersion of fish gut microbes.

### 4.3 Core genus-host phylosymbiosis and cophylogeny pattern

Phylosymbiotic relationship of host-microbiome system has been proved in vertebrates and invertebrates (43, 48, 49, 50). Indeed, significant signals of phylosymbiosis were observed in the present study (Figure 4A). In general, the basis for phylosymbiosis is that microbiome similarity among species in a community is predicted to decrease with increasing evolutionary divergence of the host organisms (51). According to current research, host-microbiota phylosymbiosis patterns may be caused by a variety of factors, including phenotypic divergence between fish that are distant in genetic relationship and co-evolution between microbiome and their hosts (52). Moreover, even patterns of host behavior or life history that may be related to phylogeny but also indirectly affect microbial communities (e.g., feeding preferences, habitat selection and morphological trait) (53). Ecological processes such as selection and drift can also shape species correlations and their associated microbial communities, leading to phylosymbiosis (42). Overall, it is important to note that phylosymbiosis is not a sign of host-microbiome adaptive co-evolution.

In the phylogenetic tree of the species collected in this study, we observed different LIPA-ASVs signals (Figure 4E). Our results showed the strongest symbiotic signal was found in Cypriniformes, while weak symbiotic signal was found in Siluriformes, Perciformes, Synbranchiales and Clupeformes. In Cypriniformes species, ASV-specific phylogenetic signal was centralized associated with Cyprinidae species. This result is in line with a few studies on vertebrate gut microbiomes (42), indicating that phylogenetic signal is strongest at finer taxonomic levels (48). In general, phylosymbiosis is mainly due to the following factors: (a) host specificity filtration, including host characteristics (e.g. immune level, metabolic system, gastrointestinal microhabitat and etc.) which is associated with the host’s phylogeny and has a remarkable influence on gut microbiota assembly, (b)t the shared evolutionary history of co-evolution between microbes and hosts (54), and (c) the above two factors often tend to work together to affect the phylosymbiosis between hosts and microorganisms (19, 51).

Our findings further showed that fish species exhibited strong signals of co-phylogeny with some of their core bacteria genera, including *Cetobacterium*, *Clostridium*_sensu_stricto_1, *Romboutsia*, *Bacteroides*, while *Aeromonas* showed no signals of co-phylogeny with their fish hosts. Still, we found that quite a few LIPA-ASVs from *Aeromonas* which is one of the dominant genera next to *Cetobacterium* and *Clostridium*_sensu_stricto_1. It has been shown that *Aeromonas* is a common pathogenic in freshwater fish and is widespread (55). These results suggested that the phylosymbiosis may not be only a partial result of co-evolution, but mainly a result of ecological or host physiological filtering. All in all, phylosymbiosis in fish species is likely to be influenced by multiple factors, e.g. combination of both ecological filtering and co-evolution of microbiota with their hosts (17). Furthermore, across the phylogeny of the sampled fish, we further observed variations in the signals of phylosymbiosis at different taxonomic levels. Cypriniformes and Perciformes species showing cophylogeny with *Cetobacterium*, while Siluriformes species did not share prevalent phylogenetic history with *Cetobacterium*. However, in the present study this could also be an artefact due to the relatively small sample size analysed in some fish orders, which may require more species to be included to more safely reveal the potential co-phylogeny signal. Specifically, different degrees of symbiosis in different fish orders had over different evolutionary timescales. After all, symbiosis is not a universal pattern of host-microbiota relationships (56, 57). For instance, existing research has reported that the gut microbes of wild fish in the Yangtze River mainly come from environmental microorganisms related to fish feeding, most of them are only ‘transient guests’ (temporary residents), and the proportion of permanent residents who form stable and adapted communities with the host is very low (58). Even though systemic symbiosis is evident at larger phylogenetic scales (different host phyla and classes) (48), this intensity can vary significantly at finer taxonomic scales (e.g. family or genus) due to differences in mechanisms and their relative strength in forming patterns (4). For example, differences in the relative contributions of factors such as diet, habitat preferences and geographical location, life history, and social interactions shape the highly variable intensity of phylosymbiosis (18, 59).

## 5 Conclusion

In the present study, we quantified the relative contribution of various factors in shaping the gut bacterial community assembly across various fish hosts in a large subtropical river of China. Our findings have demonstrated that fish host specificity is among the key drivers of gut microbiota evolution and diversification. Different taxonomic levels of host showed different degrees of contribution in variation of gut microbiota. Phylosymbiosis is evident at both global and local levels, which are jointly shaped by complex factors including ecological or host physiological filtration and evolutionary process. The core microbiota showed co-evolutionary relationships of varying degrees with different taxonomic groups. Based on our findings, we suggest that host genetic isolation or habitat variation facilitate the heterogeneous selection (deterministic process), which results in different host-core microbe specificity.

## Data availability statement

Sequencing data and relevant files have been uploaded to Sequence Read Archive with the accession number PRJNA1183623

## Author contribution

Y.L.: formal analysis, methodology, data curation, visualization, and writing (original draft); K.K: formal analysis, methodology, and writing (review and editing); H.L. and Ye. L.: data curation, methodology, resources; X.L. and J.L.: conceptualization, data curation, funding acquisition, project administration, supervision, validation, visualization.

## Supplementary materials

**Supplementary Figure 1:**
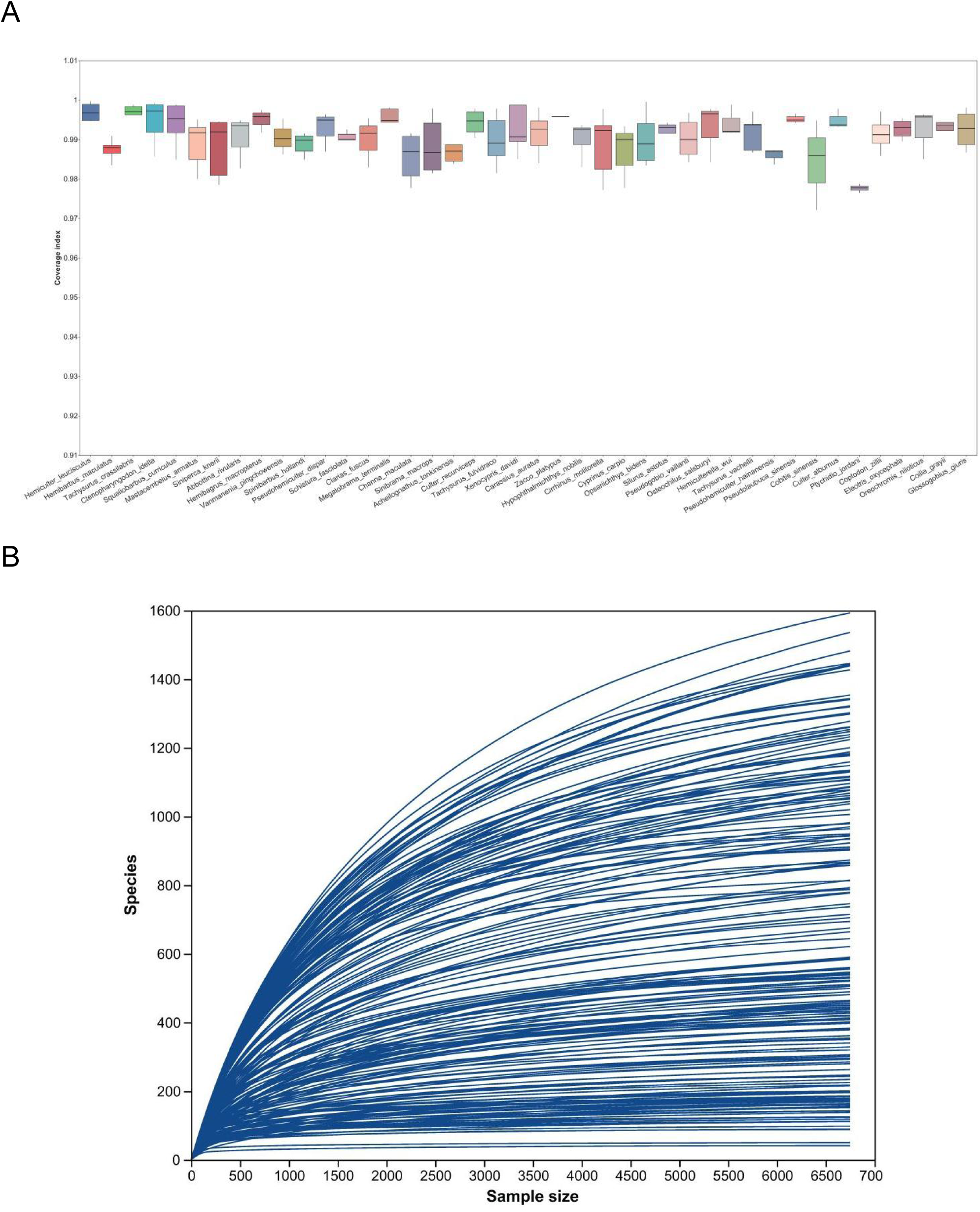
(A) Box plots of Good’s coverage of the gut microbiome samples analyzed for the fish species; (B) rarefaction curves showing the correlation between sequencing depth and bacteria taxa recovered for all gut samples sequenced.

**Supplementary Figure 2:**
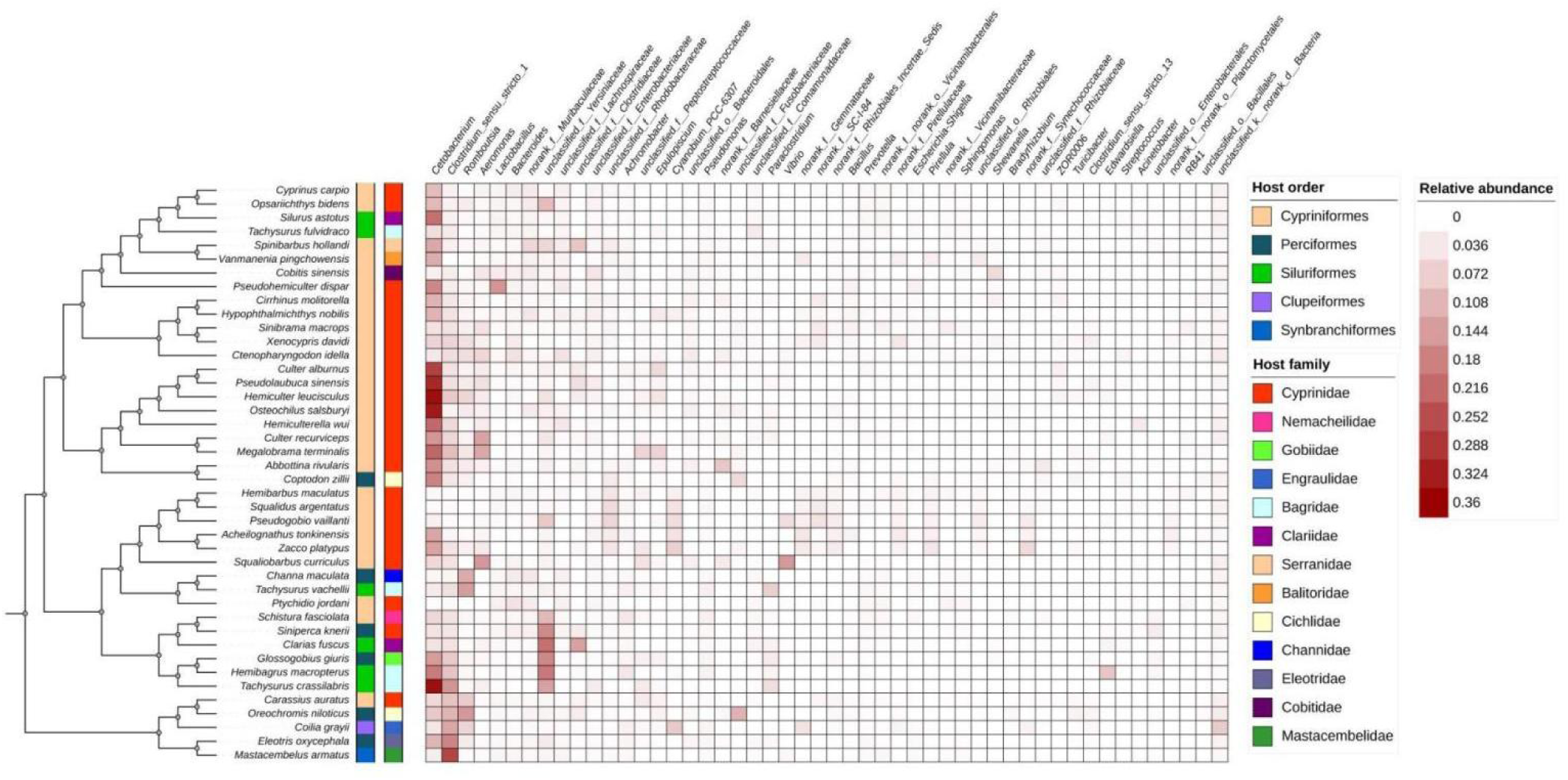
Heatmap indicated relative abundance of the top 50 bacterial genus in each of the fish species. Microbiota dendrogram according to mean of bray-curtis distances among different species

**Supplementary Figure 3:**
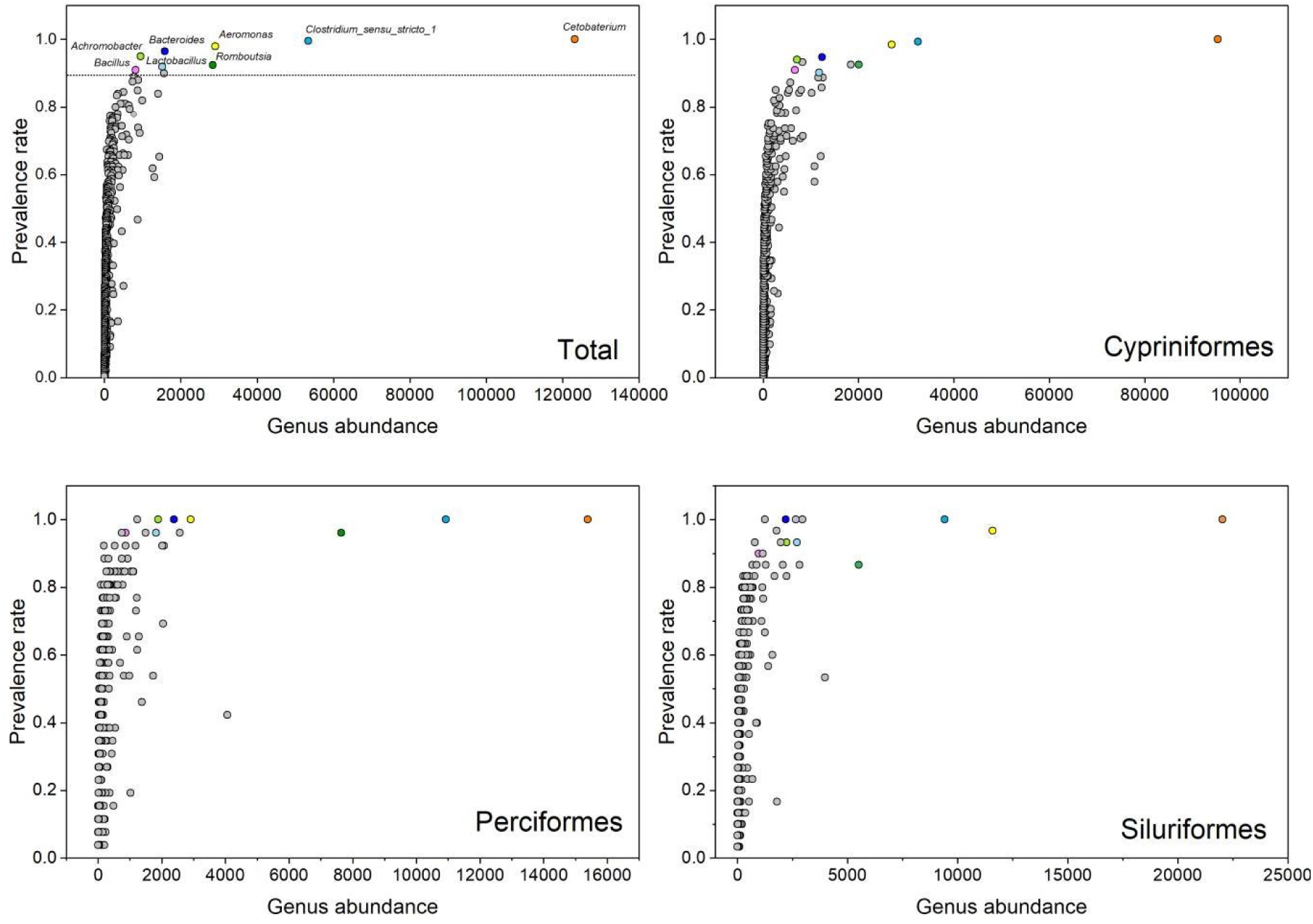
The eight mostly prevalent genera (*Cetobaterium*, *Clostridium_sensu_stricto_1*, *Aeromonas*, *Romboutsia*, *Bacteroides*, *Lactobacillus*, *Achromobacter*, *and Bacillus*) are shown in samples from three different orders (All, Cypriniformes, Perciformes, Siluriformes)

**Supplementary Figure 4:**
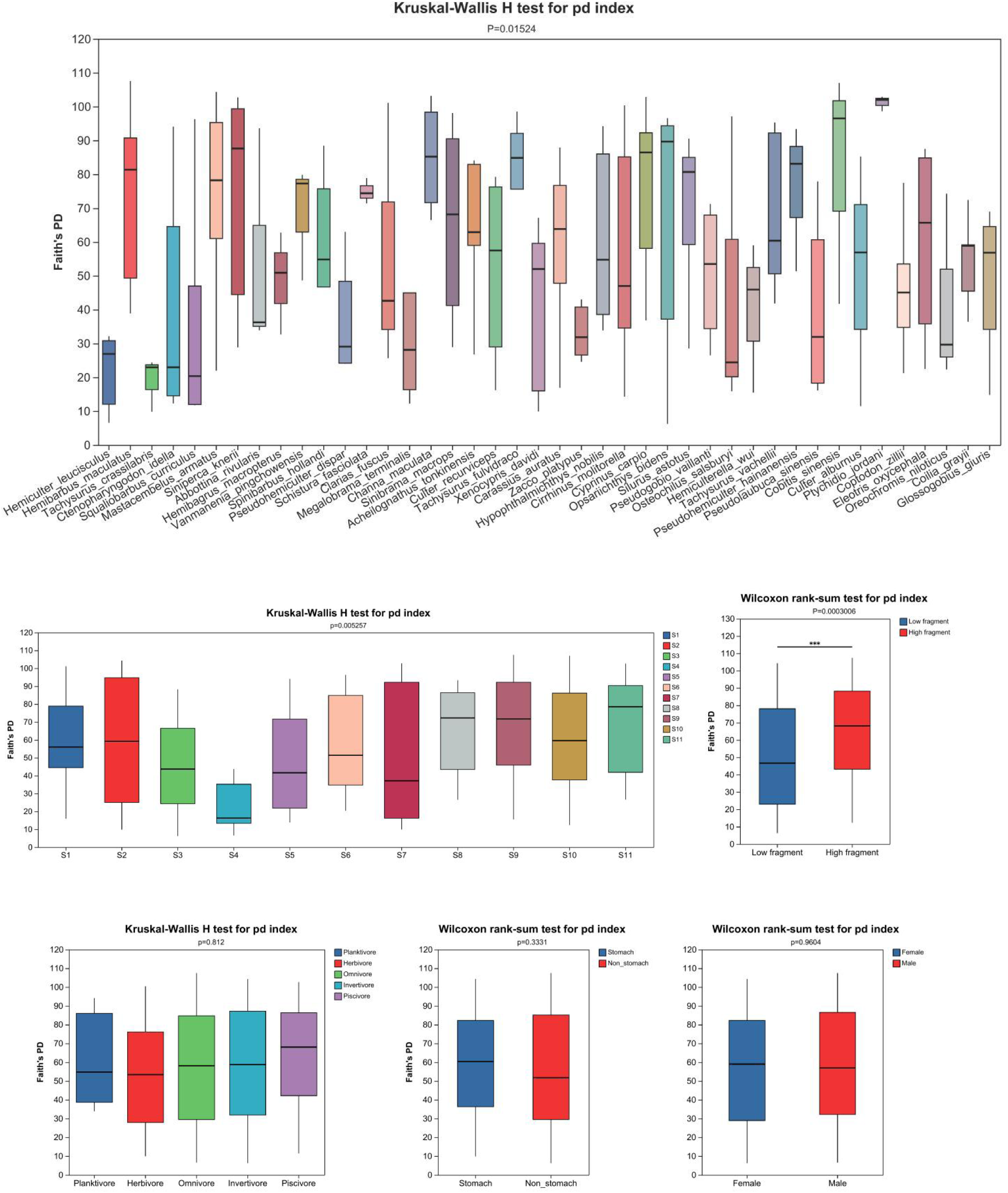
α-diversity of fish gut microbiota in hosts of different species,habitat type, geographical location, dietary group, and stomach type

**Supplementary Figure 5:**
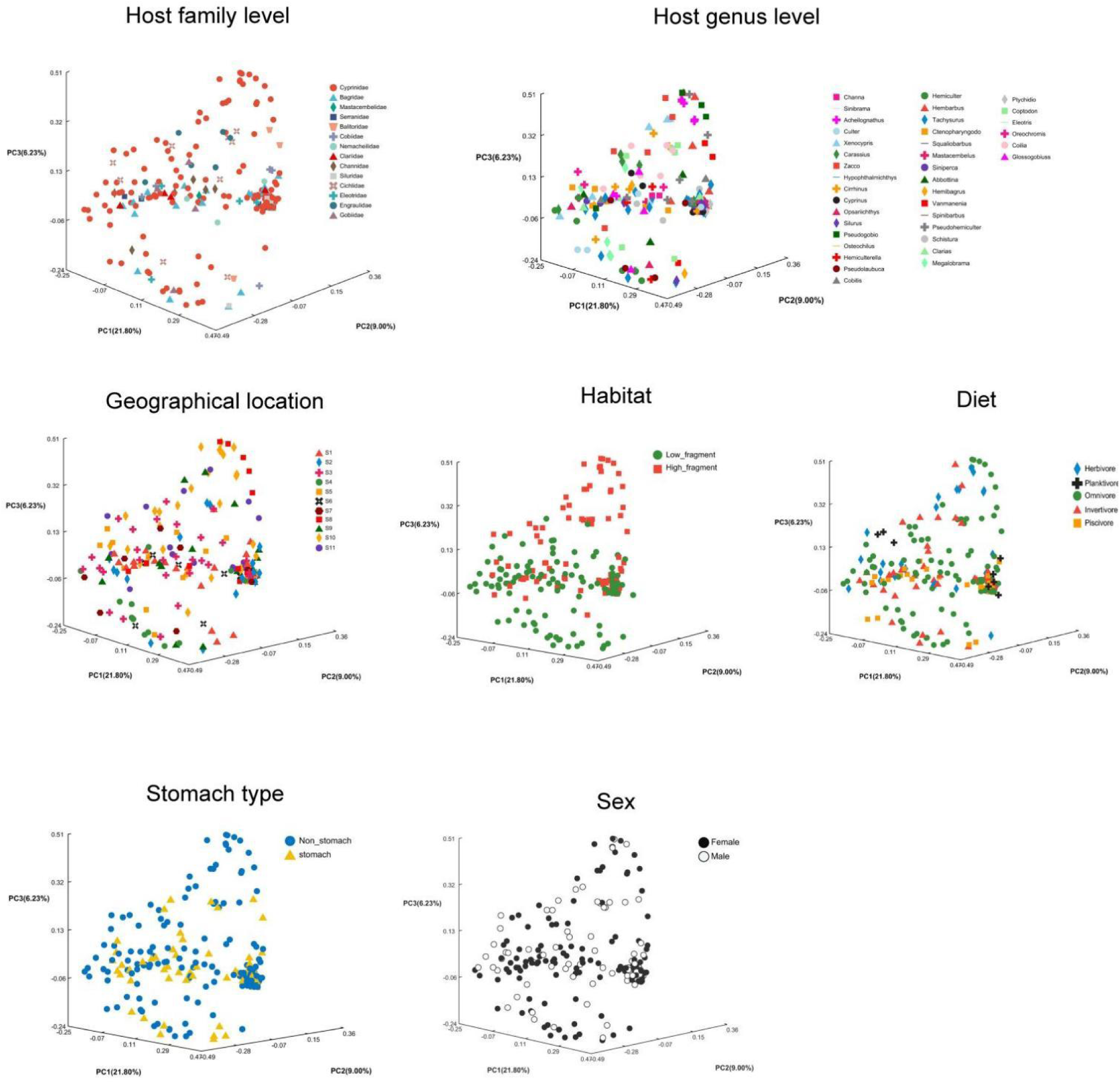
PCoA plots based on Bray-Curtis dissimilarity showing the variability in microbial composition between fish gut microbiome samples. Colour and shape of different data points corresponds to the (A) host family; (B) host genus; (C) geographical locations; (D) habitat; (E) dietary preference of host; (F) stomach type; (G) sex

**Supplementary Figure 6:**
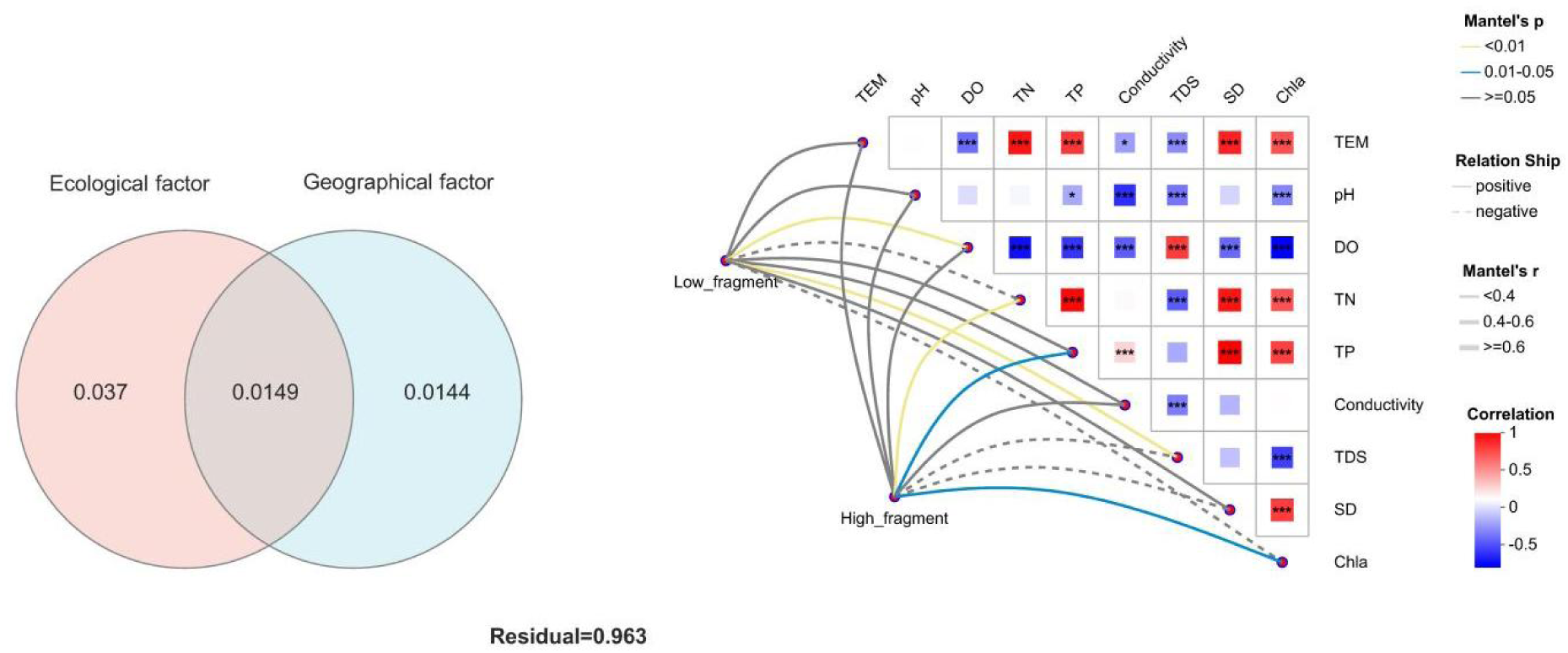
(A) Variation partitioning of environmental factors and geographical distance; (B) Relationship between environmental factors and fish gut bacterial community. Asterisks represent *P*-values: \*\*\**P* < 0.001; \*\**P* < 0.01; \**P* < 0.05.

**Supplementary Figure 7:**
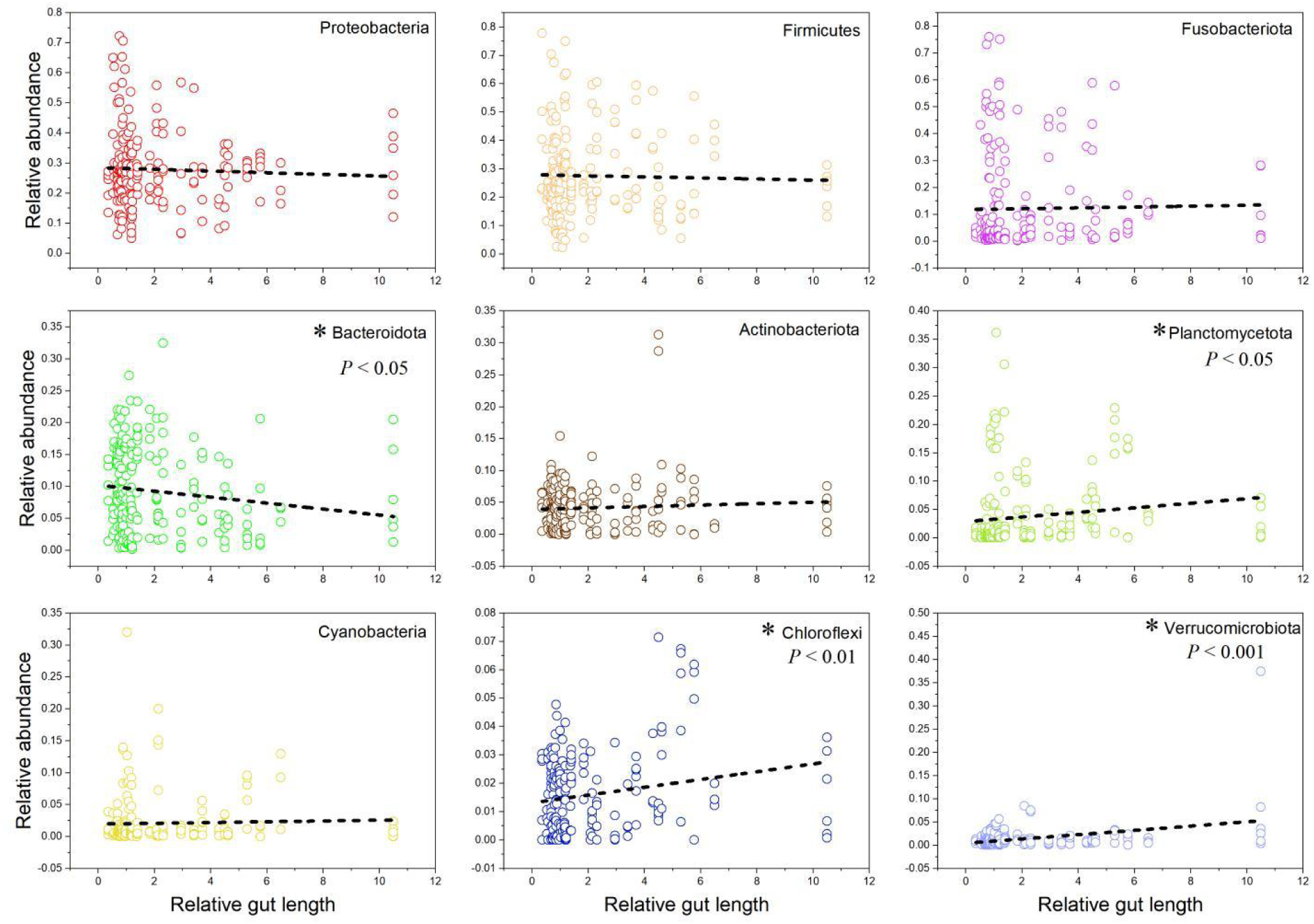
Linear regression of relative abundance of gut core microbiome (the phylum level) among all fish species vs. their relative gut length, (A) Proteobacteria; (B) Firmicutes; (C) Fusobacteriota; (D) Bacteroidota; (E) Actinobacteriota; (F) Planctomycetota; (G) Cyanobacteria; (H) Chloroflexi; (I) Verrucomicrobiota.

**Supplementary Figure 8:**
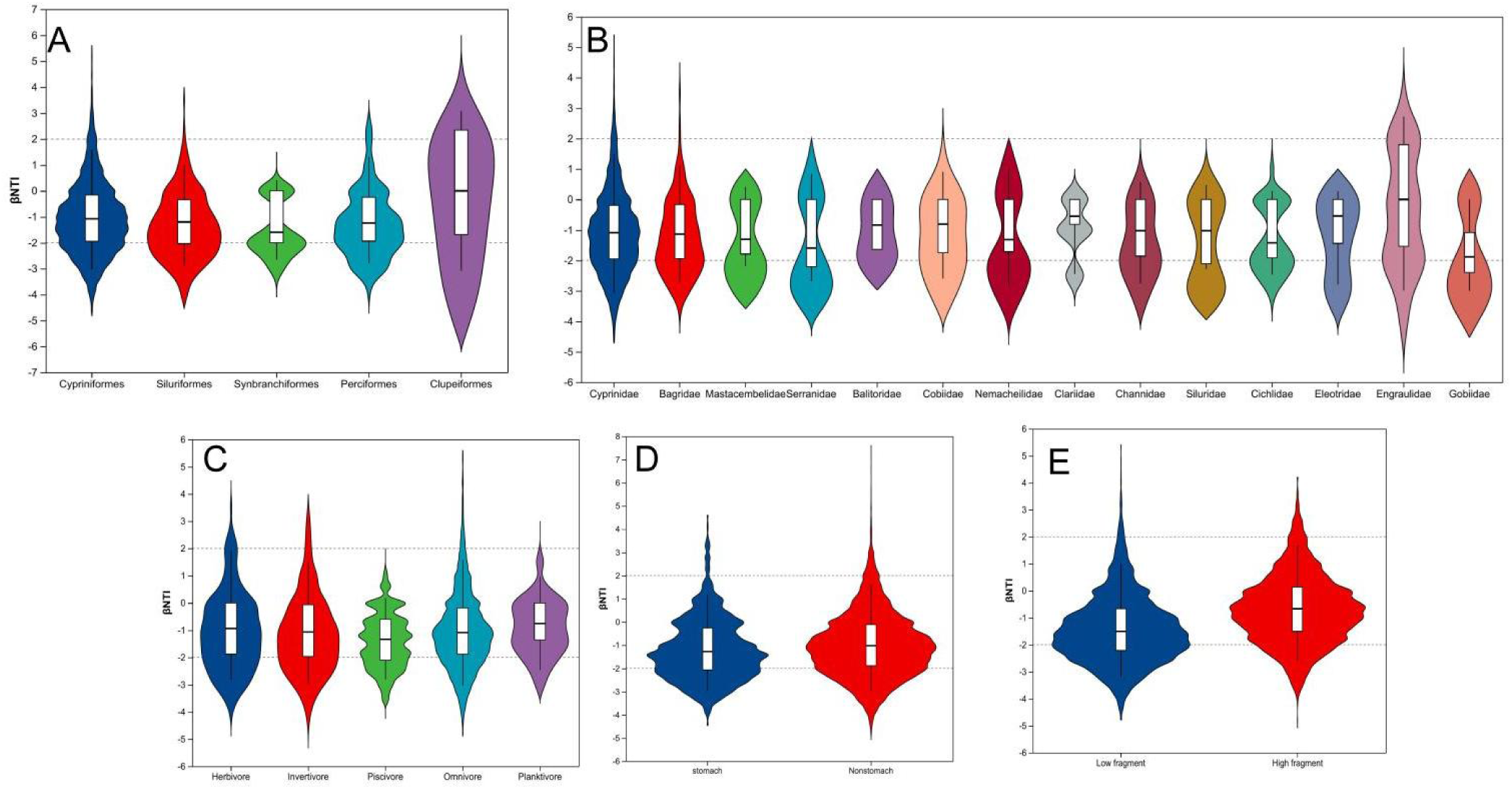
Null model was used to analyze the assembly process of gut microbial community in hosts of different species (A,B), dietary group (C), stomach type (D) and habitat type (E).

Supplementary Table 1 Metadata of the ecological and sampling information of the fish species for statistical analysis.

(Supplementary files. Table S1.xlsx)

**Supplementary Table 2.** The variance of fish gut microbiota composition explained by environmental factors in the lower research of the Pearl River.

| Environment factor | $R^2$ | $P$-value |
| --- | --- | --- |
| TEM | 0.0864 | 0.001 |
| pH | 0.0188 | 0.172 |
| DO | 0.1629 | 0.001 |
| TN | 0.1822 | 0.001 |
| TP | 0.1337 | 0.001 |
| Conductivity | 0.0022 | 0.795 |
| TDS | 0.1357 | 0.001 |
| SD | 0.1088 | 0.001 |
| Chla | 0.166 | 0.001 |

